# Sex differences in senescence burden within human osteoarthritic synovial fibroblasts

**DOI:** 10.64898/2026.08.14.744916

**Authors:** Garrett Sessions, Tarek Zikry, Lauryn E. Bailey, Jacqueline Shine, Richard Loeser, Sam Wolff, Jeremy Purvis, Brian Diekman

**Affiliations:** Department of Cell Biology and Physiology, The University of North Carolina at Chapel Hill, Chapel Hill, NC 27599, Unites States of America; School of Data and Information Sciences, The University of North Carolina at Chapel Hill, Chapel Hill, NC 27599, United States of America; Computational Medicine Program, The University of North Carolina at Chapel Hill, Chapel Hill, NC 27599, United States of America; Lineberger Comprehensive Cancer Center, University of North Carolina at Chapel Hill, North Carolina 27599, United States of America; Department of Genetics, The University of North Carolina at Chapel Hill, Chapel Hill, NC 27599, United States of America; Lampe Joint Department of Biomedical Engineering, University of North Carolina at Chapel Hill, Chapel Hill, North Carolina 27599 and North Carolina State University, Raleigh, North Carolina 27695, United States of America; Thurston Arthritis Research Center, The University of North Carolina at Chapel Hill, Chapel Hill, NC 27599, United States of America; Division of Rheumatology, Allergy, & Immunology, The University of North Carolina School of Medicine, Chapel Hill, NC 27599, United States of America

**Author notes:** **Corresponding authors:** Jeremy Purvis, Department of Genetics, The University of North Carolina School of Medicine, 11018C Mary Ellen Jones, Campus Box #7264, Chapel Hill, NC 27599-7264, Brian Diekman, Department of Biomedical Engineering, The University of North Carolina School of Medicine, 4111-A Thurston, Campus Box #7280, Chapel Hill, NC 27599.

## Abstract

**Objective:** Cellular senescence has been shown to underlie many age-related diseases, including osteoarthritis (OA). In addition to age, biological sex is an OA risk factor with females at greater risk of hand and knee OA. We profiled the senescence burden in OA human synovial fibroblasts while accounting for these factors to understand how senescence may contribute to the increased burden of OA in females.

**Methods:** Synovial fibroblasts were isolated from tissue obtained at knee arthroplasty for OA from 10 male and 10 female donors. Single cell multiplexed immunofluorescence imaging was used to profile the senescence burden in samples age-matched to account for the differences in chronological age. Clustering was performed using stability and generalizability scoring.

**Results:** Independent of chronological age, OA synovial fibroblasts from female donors showed higher levels of senescence associated proteins p16, p21, p53, phospho-p65, IL-6, and IL-8. Assessment of oxidative stress associated proteins NRF2, SEPP1, NQO1 and TXNIP indicated a lower capacity for female cells to respond to oxidative stress. Clustering analysis revealed male and female enriched clusters. The female-enriched clusters showed higher levels of senescence-associated proteins and an increased oxidative stress response.

**Conclusions:** OA synovial fibroblasts from female donors demonstrated higher levels of senescence associated markers, lower ability to respond to oxidative stress, and increased senescence with increasing age. These findings indicate that female synovial fibroblasts are more likely to show markers of senescence and oxidative stress, suggesting senescence can contribute to the increased incidence of osteoarthritis in women.

## Introduction

Osteoarthritis (OA) is a leading cause of pain and disability in older adults, with 30.8 million US adults estimated to suffer from OA (1). Despite extensive investigation, treatment for OA is largely limited to pain management until total joint replacement is required. Aging and female sex are two of the strongest risk factors for OA, with the excess cases in female prevalence increasing with age (2). One potential explanation for this difference is the accumulation of senescent cells in the joint. Senescence emerges in response to persistent stress and synovial fibroblasts adopt features consistent with the senescence phenotype(3,4). Previous work has drawn connections between senescence burden in the joint space and osteoarthritis progression(3,4), but has not considered interactions between biological sex and cellular senescence.

To account for the possibility of age- and sex-related effects on synovial fibroblast senescence, we collected pairs of synovial tissue from male and female patients undergoing total knee arthroplasty for OA. This strategy allowed us to study the effects of biological sex and chronological age both independently and as interacting factors.

Previous studies examining gene expression demonstrated an age effect for many senescence-associated transcripts (5,6). However, aging exacerbates the decoupling of transcript levels and protein levels, which may compromise the ability to interpret senescence features based on gene expression alone (7). Further, transcriptomic measurements are limited in their ability to resolve key cellular processes like protein localization and phosphorylation state, which are critical to the regulation of senescence and the senescence associated secretory phenotype (SASP). A final concern is that bulk analysis can obscure distinct subpopulations of senescent cells, known as senotypes (8). To address these challenges, we took advantage of the single-cell protein profiling through iterative indirect immunofluorescence imaging (4i) (9).

Our work interrogates a heterogenous population of cells and identifies critical senescent subpopulations that may contribute to the observed differences in the progression of age-related diseases. Here we revealed the elevated expression of canonical senescence markers in female OA synovial fibroblasts compared to males across the spectrum of chronological age. We leveraged 4i as a novel method of interrogating primary human synovial fibroblasts and resolved specific subpopulations of synovial fibroblasts that drive the distinct features of senescence in females. By understanding how the senescence burden differentially accumulates between sexes, the field will be better equipped to target senescence for OA therapy.

## Materials and Methods

### Tissue procurement and culture conditions

Primary human synovial fibroblasts were isolated from synovial tissue collected during total knee joint arthroplasty for OA as described (10). The use of de-identified human tissues was approved by the University of North Carolina Institutional Review Board. Frozen cells from male and female donors were thawed and plated in parallel for use in experiments. Cultures were confirmed to be enriched for synovial fibroblasts by flow cytometry for CD73 and CD105 (Supplemental Figure 1). See Supplemental Methods for culture details.

### Iterative Indirect Immunofluorescence Imaging (4i)

Sample preparation followed the original 4i protocol (9). Testing of primary antibodies (Supplemental Table 1) was performed to ensure correct staining above background and elution using established 4i protocols. See Supplemental Methods for additional details.

### Statistical Analysis

Statistical analysis was performed using GraphPad Prism 11 and included the use of Welch’s t test and simple linear regression. Z-normalization was performed on a per feature basis. The predictive modeling of sex and age was performed using resources from the Python package sklearn. Sex was evaluated using a linear logistic regression model and a random forest model. Age was evaluated using the linear ElasticNet model and a random forest model. Clustering was determined through a combination of literature expectations, stability scoring, and generalizability scoring using tools from the sklearn package. All models were evaluated using a standard 80/20 train/test split. Fisher’s Exact Test was used to calculate adjusted p-values for the clustering analysis.

## Results

Primary human OA synovial fibroblasts were isolated from ten pairs of age-matched male and female donors, which allowed us to investigate the complex relationship between biological sex and chronological age (Supplemental Table 2). Utilizing 4i (9), single-cell proteomics data were collected across five 96 well plates that each contained the entire donor set, with common markers measured across plates to allow merging of replicates (Supplemental Table 3). Markers were selected to provide insight into the cellular senescence burden, SASP production, and oxidative stress load on individual cells.

When pooled by biological sex, female donors showed higher expression of canonical senescence markers such as p16 (p=0.0016) and p21 (p<0.0001) but did not show a difference in p53 (p=0.8262). Female donors also showed an increase in SASP associated markers phospho-p65 (p<0.0001), IL-8 (p=0.0017) and IL-6 (p<0.0001) (Figure 1a). We also investigated nuclear p16 concentration in age matched donor pairs. For all age matched donor pairs except one (71-year-old female vs. 73-year-old male), the female donor in the pair showed significantly higher p16 expression than the male donor (Figure 1B). The levels of nuclear p16 showed a modest age-related increase in both male and female donors as assessed by the slope of linear regression analysis, which was significantly higher in female donors (p<0.0001) (Figure 1C). Female donors also showed a moderately higher age-related increase in p21, whereas the slope for male donors was not significantly different from zero (Figure 1C). Female donors showed increased phospho-p65 with age, but male donors demonstrated a slight decrease in phospho-p65 with age (Figure 1C). Male and female donors showed an equivalent age-related increase in p53, and approximations of Interleukin-6 (IL-6) and Interleukin-8 (IL-8) secretion using cytoplasmic concentrations showed no increase or a limited decrease in levels with age in both males and females.

We next quantified markers of oxidative stress, which has previously been associated with osteoarthritis (11). Nuclear factor erythroid 2-related factor 2 (NRF2) is a master regulator of antioxidant genes and is increased under conditions of oxidative stress (12). The nuclear concentration of NRF2 was higher in females compared to males (Figure 2A). Thioredoxin-interacting protein (TXNIP), which is a known pro-oxidative stress protein, was higher in females compared to males. The anti-oxidant protein Peroxiredoxin 6 (PRDX6) was also higher in females, suggesting a potential compensatory response. However, other markers of a robust antioxidant response were actually lower in female donors. Despite being a canonical target of NRF2, NQO1 was reduced in female synovial fibroblasts. Selenoprotein P (encoded by SEPP1/SELENOP) provides selenium to antioxidant proteins, and this protein was also reduced in females when all ages were combined. By breaking the donors out by age-matched pairs, we observed the female/male pairs under the age of 64 show higher levels of SEPP1 in females, while pairs over the age of 65 show higher SEPP1 levels in males (Figure 2B). When chronological age is taken into consideration, SEPP1 shows a significant age-related decline in female donors (Figure 2C). This is distinct from male donors, which do not show any significant slope (p=0.1161). Reduced SEPP1 in serum has been shown to correlate with reduced scores on the Functional Ability Questionnaire in patients with OA, suggesting a possible role for SEPP1 in mitigating disease severity (13). Similar, though less pronounced, trends are observed in the levels of NRF2 and NQO1, with both markers showing a greater age-related decline in females compared to males (Figure 2C). TXNIP showed indistinguishable rates of change due to chronological age between female and male donors, with females maintaining a higher concentration regardless of age. Together, these data indicate that synovial fibroblasts from females likely exhibit a higher level of oxidative stress as compared to males but a reduced baseline capacity for resolving it, with a sharper decline in antioxidant capacity with age.

**Figure 1.**
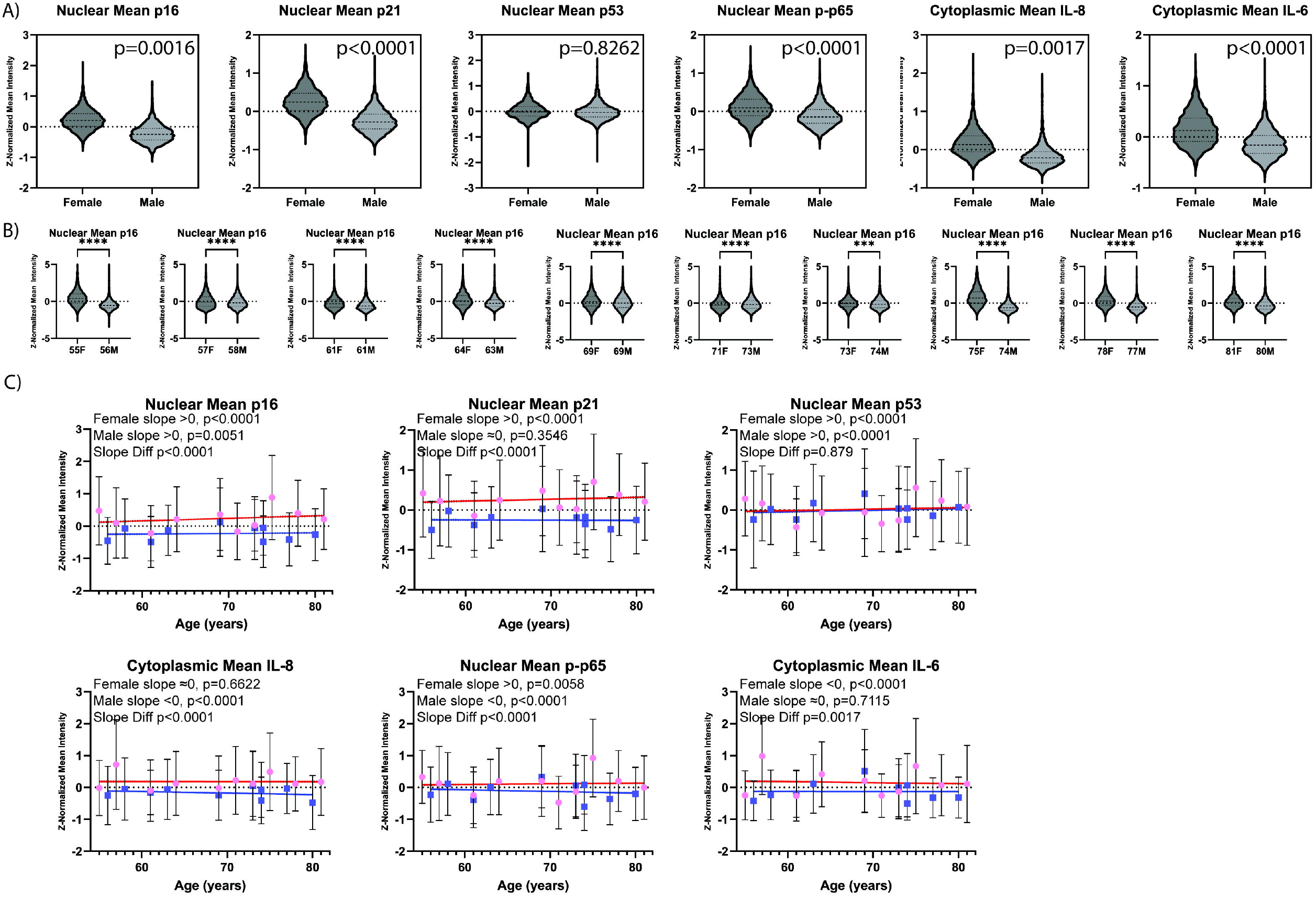
Levels of canonical senescence markers. A) The levels of canonical senescence markers p16, p21, p-p65, IL-8 and IL-6 are all elevated in female OA synovial fibroblasts compared to male. B) Direct comparison of p16 in matched pairs of donors shows elevated levels of p16 in females at all paired ages. C) Linear regression analysis was performed for each canonical senescence marker with respect to chronological age. The directionality of slope was derived using the 95% confidence interval and the statistical significance of the difference between slopes is shown in the inset table.

**Figure 2.**
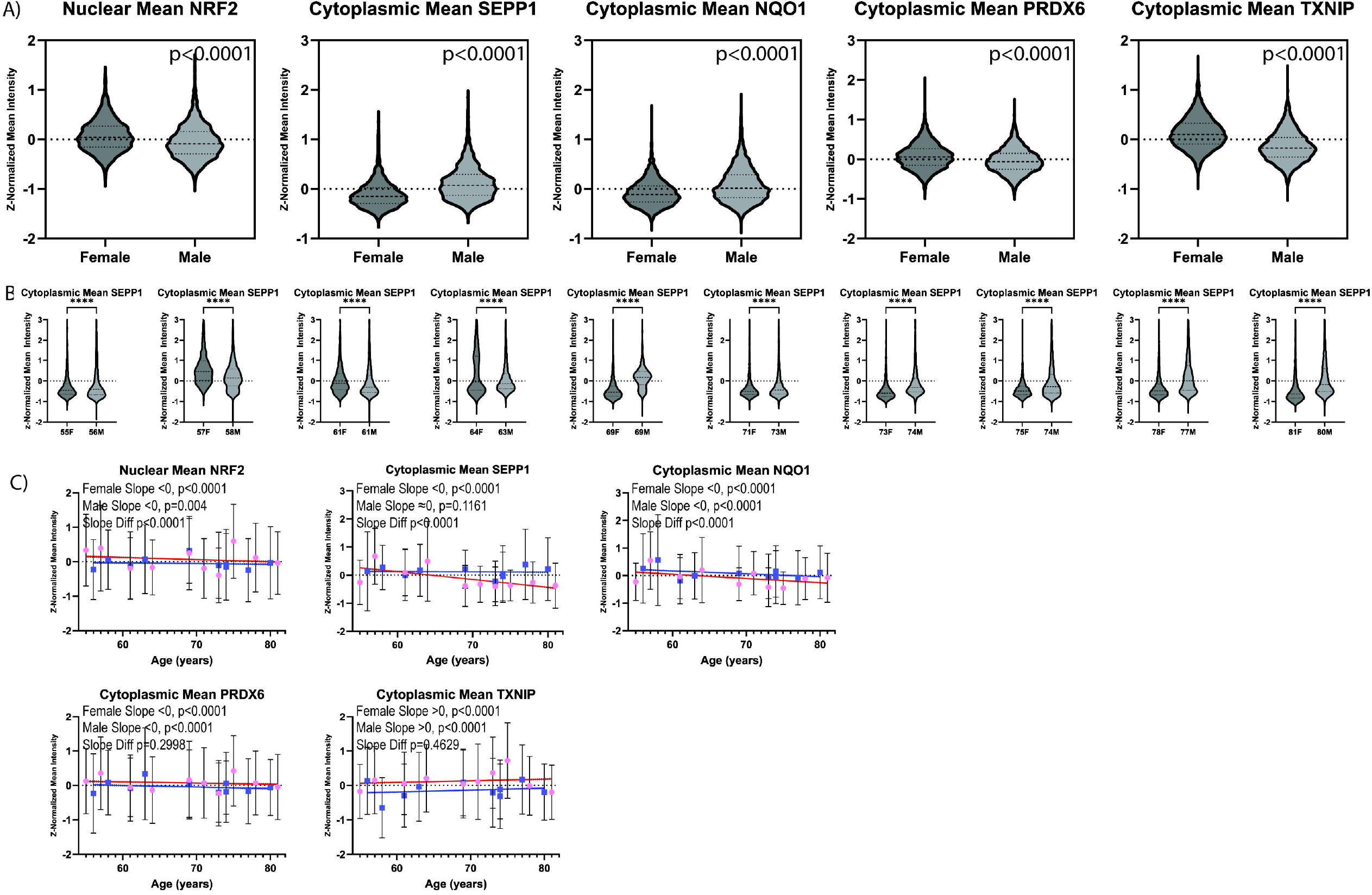
Levels of selected markers of oxidative stress. A) The levels of NRF2, TXNIP and PRDX6 are elevated in female OA synovial fibroblasts compared to male. SEPP1 and NQ01 are both reduced in female OA synovial fibroblasts compared to male. B) Direct comparison of SEPP1 in matched pairs of donors shows elevated expression of SEPP1 until age 64 and reduced levels of SEPP1 in after age 64 in female donors compared to males. C) Linear regression analysis was performed for each oxidative stress marker with respect to chronological age. The directionality of slope was derived using the 95% confidence interval and the statistical significance of the difference between slopes is reported in the inset table.

These analyses indicate that OA synovial fibroblasts harbor a complex assortment of molecular markers that may be responsible for age- or sex-related differences in disease progression. Thus, we next used predictive modeling to ask what combination of markers could best separate samples by age and sex (Supplemental Table 4). The logistic regression sex models displayed strong predictive performance (Supplemental Figure 2) and age was less predictive than sex, which supports the idea that these donors may represent a similar biological age at the time of joint replacement despite a wide range of chronological age (Supplemental Figure 3). We also explored the possibility that certain samples contained discrete clusters of cells with multi-factor senescent features. We used the Model Explorer method (14) to choose the optimal number of clusters with subsample proportion of 0.5 after taking a random sample of 100 cells per plate for computational efficiency over 500 iterations with adjusted rand index (ARI) as the metric of comparison. We further assessed the predictability of clusters with a random forest of 500 trees on an 80/20 train test split. By projecting these clusters onto a PHATE map, we demonstrated that clusters from each plate produce a similar structure and that the clusters map in similar ways across all plates (Supplemental Figure 4A). We observed how individual cells were distributed throughout these clusters (Supplemental Figure 4B) and projected the measured features for each cluster as a heatmap (Supplemental Figure 4C) to validate the cluster unification strategy.

Quantification of the difference between canonical senescence markers within clusters shows two key clusters, which we refer to as the High Concentration Senescent cluster and the Stereotypical Senescence cluster. These two clusters both have elevated concentrations of p16, p21, and p53 and differ mainly in their measured nuclear area (Figure 3A). The High Concentration Senescent cluster and the Stereotypical Senescence cluster also have higher concentrations of phospho-p65, a known regulator of the SASP, and cytoplasmic concentrations of IL-6 and IL-8, used here as an approximation of SASP secretion (Figure 3B). Both clusters display a robust response to oxidative stress, showing increased NRF2 and the antioxidative enzymes SEPP1, PRDX6, and NQO1, as well as increased levels of the pro-oxidative stress marker TXNIP (Figure 3C). A third senescent population, the Large Nuclear Senescence cluster, has lower concentrations of canonical senescent markers, but the large nucleus results in elevated levels of total nuclear p21, p53, and p-p65 compared to the High Concentration Senescence cluster (Supplemental Figure 5). This split between a high concentration and high total protein subpopulation aligns with previous findings from long-term senescence induction experiments (15). The non-senescent cluster has feature measurements more characteristic of healthy synovial fibroblasts, with lower concentrations of stress and senescence associated markers. The Stereotypical Senescence cluster has very few cells present regardless of sex or age but also represents the strongest senescence phenotype, with the smallest variance in the levels of canonical senescence features. Both the High Concentration and Stereotypical Senescence clusters have a statistically significant enrichment in cells from female donors (Figure 3D). Given that these two clusters represent the highest concentration of senescence and SASP markers, they could be responsible for the observed differences in bulk senescence burden accumulation between sexes.

**Figure 3.**
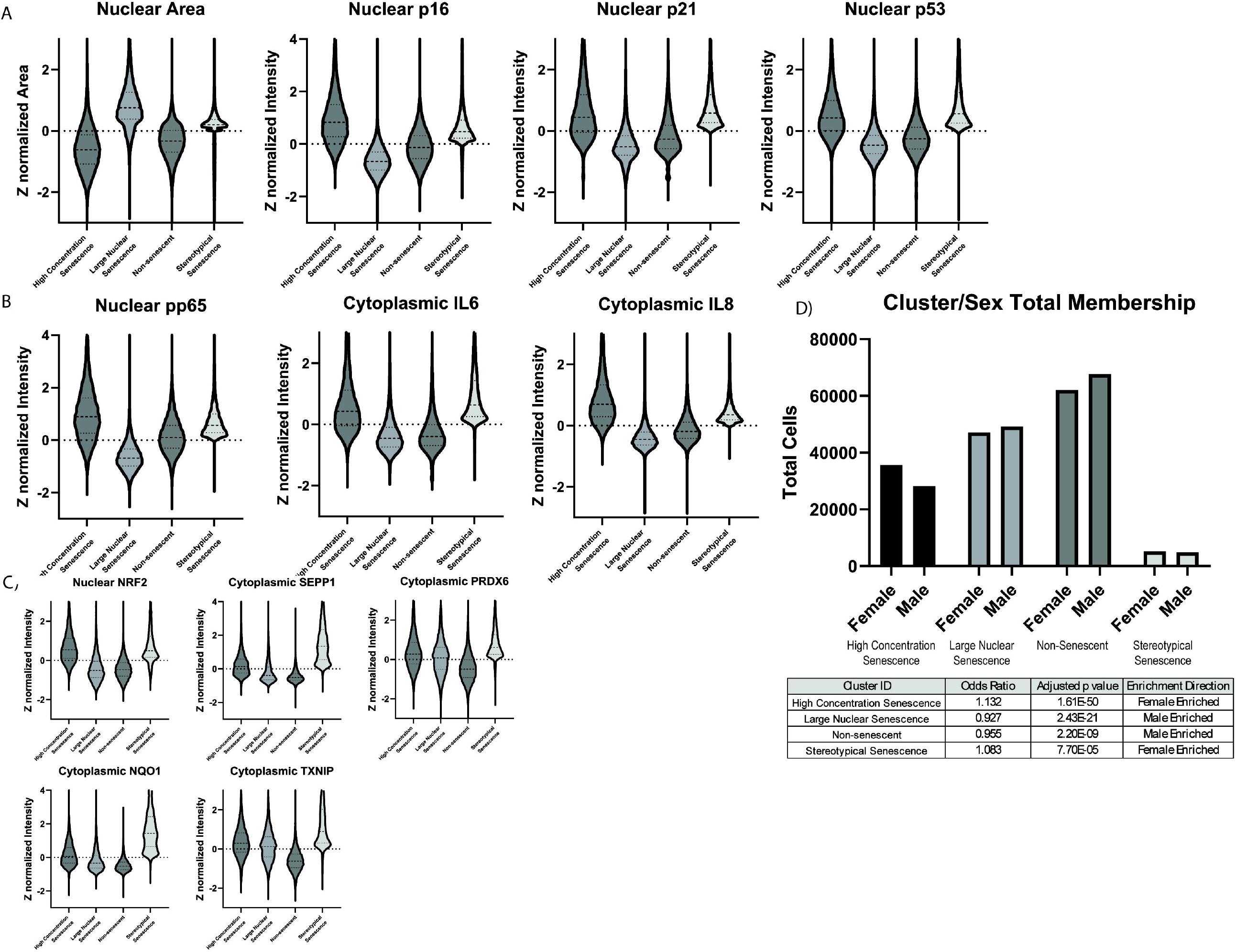
Classification of clusters and between clusters dynamics. A) The distributions of canonical senescence features nuclear area, p16, p21, and p53 are shown between unified clusters. B) The distribution of p-p65 and SASP factors IL-6 and IL-8 are shown between unified clusters. C) The distribution of markers of oxidative stress are shown between unified clusters. D) The sex bias between unified clusters and total number of cells per unified cluster is plotted. The results of a Fisher’s Exact Test are shown below with adjusted p value for each cluster and an enrichment direction derived from the odds ratio.

## Discussion

The progression of age-related diseases, such as osteoarthritis, is highly heterogeneous with numerous modulating factors including genetic background, prior injury, comorbidities, and environmental exposure. Despite the challenges inherent to studies involving primary human cells, strong evidence emerged from this work to support differences in the underlying senescence burden between male and female donors. Although we observed modest age-related differences in a small number of key senescence markers, the dominant factor predicting protein concentration was biological sex. Cells from female donors had higher levels of senescence associated proteins such as p16 and p21, as well as increased concentrations of SASP factors such as IL-6 and IL-8 (Figure 1A). This suggests that the burden of senescent cells may play a different role in the progression of OA in females, which could inform the design of future clinical trials testing senolytics as therapies to slow OA progression.

While sex, not age, was the primary driver of differences in senescence markers, the concentrations of certain canonical senescence markers diverged with age depending on biological sex. For example, only the females demonstrated an age-related increase in the nuclear concentration of p21, with the male donors demonstrating a slope that was not significantly different from zero (95% CI -0.002 to 0.0006). The regulation of SASP proteins also differed between the sexes as a function of age. Phospho-p65, a critical regulator of the NFKB pathway and SASP production, increased with age in females but decreased with age in males. This difference in upstream regulators was reflected in the cytoplasmic levels of proinflammatory SASP factors such as IL-8, which decreased with age in males (Figure 1C).

Oxidative stress can initiate and sustain cellular senescence (15). We noted several interesting interactions between sex and age for oxidative stress response proteins that may help to explain the differential disease burden. SEPP1, despite being higher in females at younger ages, declined with age in female donors but not males, ultimately resulting in a significantly lower concentration of SEPP1 with advanced age. NRF2 was always present at a higher concentration in female donors but did have a steeper decline with age in females compared to males (Figure 2C). The sharper decline in the ability to resolve oxidative stress implies an earlier susceptibility to senescence induction and to the production of a pro-inflammatory SASP, potentially driving joint damage and age-related OA.

In our cluster analysis, the High Concentration Senescence cluster was of particular interest because it was highly enriched (p=1.61e-50) for female cells. In addition to canonical senescence markers p16, p21, and p53, this cluster also contained cells with higher concentrations of phospho-p65 and both IL-6 and IL-8, indicating a strong SASP response within this subpopulation. These cells are also undergoing the strongest response to oxidative stress, with elevated NRF2 and moderately higher SEPP1, NQO1, and PRDX6. These features are indicative of cells that are attempting to resolve elevated oxidative stress. TXNIP is also elevated in the High Concentration Senescence cluster, suggesting that these cells are under significant oxidative stress and have persistently failed to resolve it. The Stereotypical Senescence cluster was similarly enriched for female cells (p=7.7e-5) and showed high concentrations of senescence markers as well as an increased nuclear area while making up only 3.41% of all cells. This percentage aligns with prior research regarding the stereotypical burden of senescent cells, and these cells may contribute to the features commonly identified in bulk assays.

In conclusion, synovial fibroblasts isolated from females that underwent total joint replacement for OA have a greater burden of senescent cells compared to males. This increased senescence burden is characterized by elevated levels of the canonical markers p16 and p21, as well as increased concentration of SASP factors such as phospho-p65, IL-6 and IL-8. With aging, female cells also display a reduced capacity to resolve oxidative stress. Clustering cells according to single-cell proteomic profiles revealed the presence of two senescence clusters that are enriched for female cells. Both the High Concentration Senescence and Stereotypical Senescence clusters have elevated p16, p21, p53 and increased SASP factors phospho-p65, IL-6 and IL-8. These subpopulations of senescent cells may contribute to the differential disease burden in OA seen at the population level.

