## Supplemental Methods for "Sex differences in senescence burden within human osteoarthritic synovial fibroblasts"

**Supplemental Materials and Methods**

**Tissue procurement and culture conditions**

Synovial cells were passaged twice and cultured at 37 °C and 5% CO_2_ in DMEM (Glibco, 11995-065) with 10% fetal bovine serum (FBS; Sigma, TMS-013-B), GlutaMAX (Glibco, 35050-061), and penicillin/streptomycin (P/S; ThermoFisher Scientific 15140148) and preserved in liquid nitrogen.

**Iterative Indirect Immunofluorescence Imaging (4i)**

Hoechst (Sigma-Aldrich 33258) at 400 ng/ml was used as a nuclear stain. Secondary antibodies were Alexa Fluor 647 Donkey anti-goat (Invitrogen A21447), Alexa Fluor 568 Donkey anti-mouse (Invitrogen A10037), and Alexa Fluor 568 Donkey anti-rabbit (Invitrogen A10042).

Stitched 5x5 images were collected using a Nikon Ti2 Eclipse inverted microscope using a Plan Apo LambdaD 20x objective lens (NA=0.8) with a Teledyne Photometrics Kinetix sCMOS camera. Image acquisition performed using the following filer cubes: DAPI (Semrock DAPI-3060A), AF488 (Semrock GFP-4050B), AF568 (Semrock mCherry-C), AF647 (Semrock LED-Cy5-5070A). Image acquisition and post-processing was performed using NIS-Elements HCA JOBS software. Background corrected images from each round of 4i were aligned using the StackReg library and manually confirmed. Segmentation was performed using CellPose and the region properties library of Scikit-image were used for feature quantification and extraction.
