## Supplementary figures and images for "Sex differences in senescence burden within human osteoarthritic synovial fibroblasts"

### Supplemental Figure 1

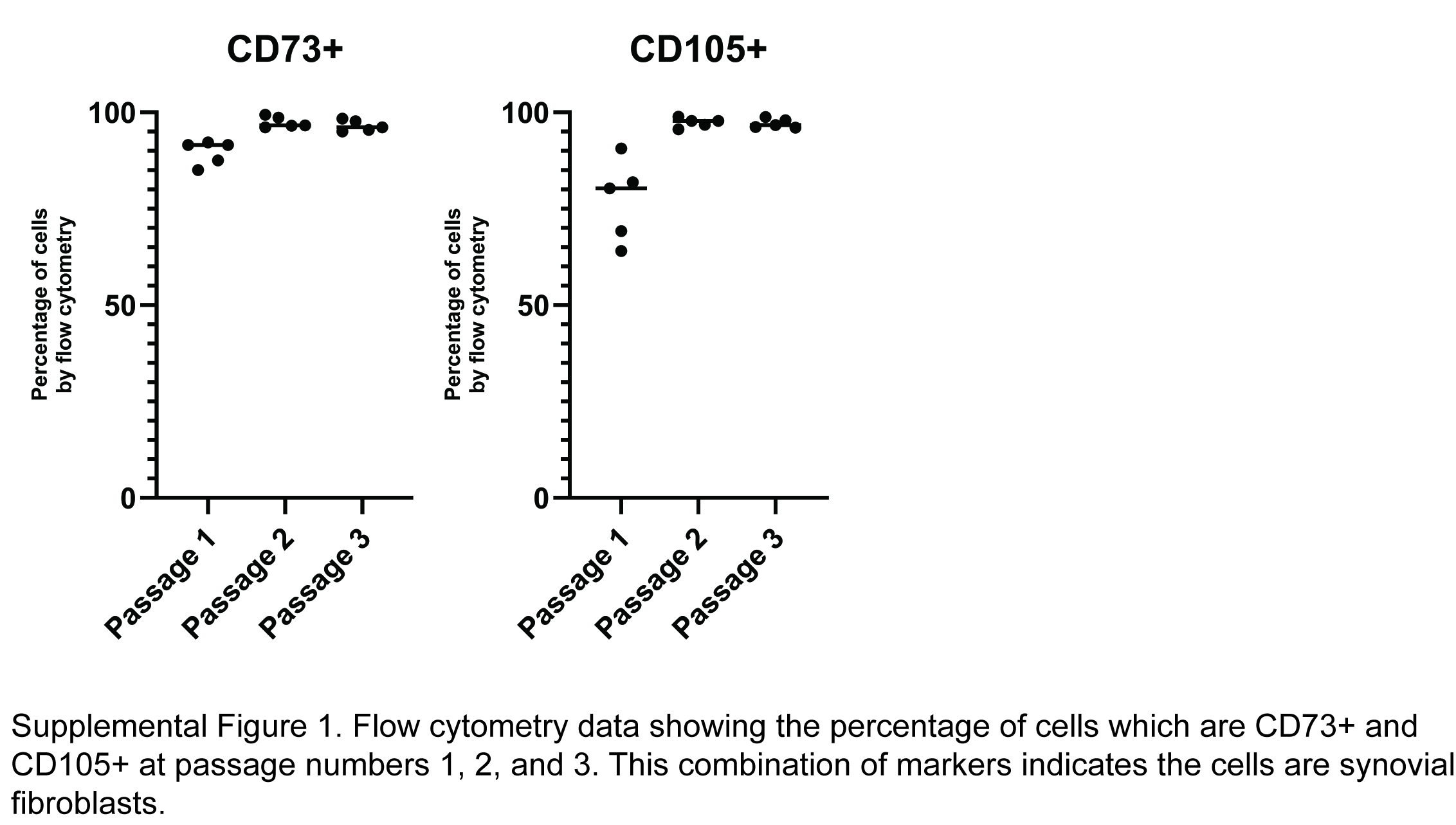

### Supplemental Figure 2

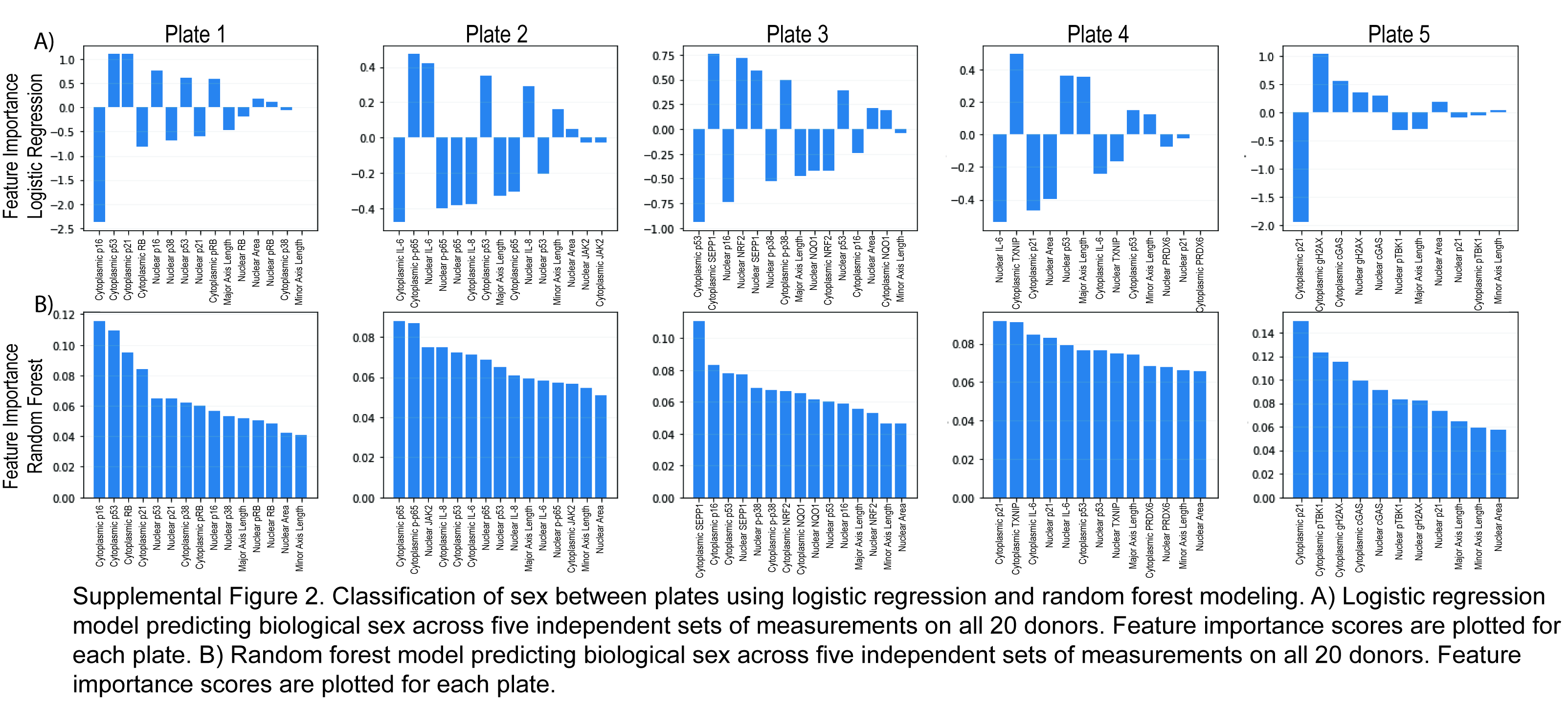

### Supplemental Figure 3

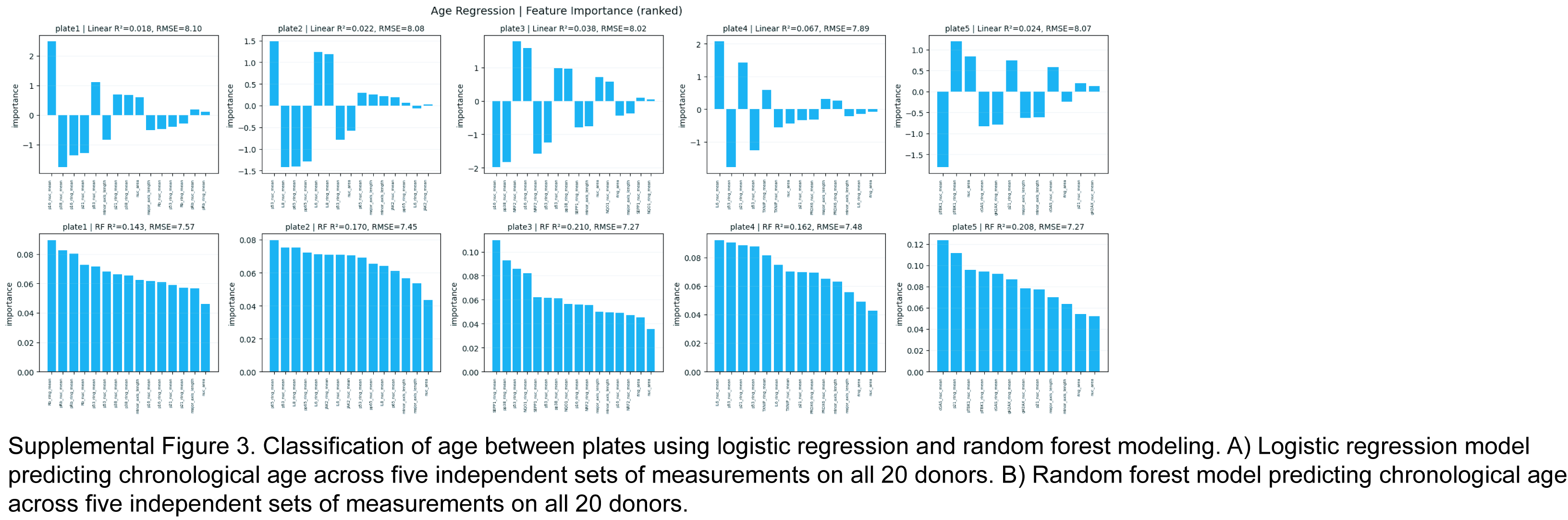

### Supplemental Figure 4

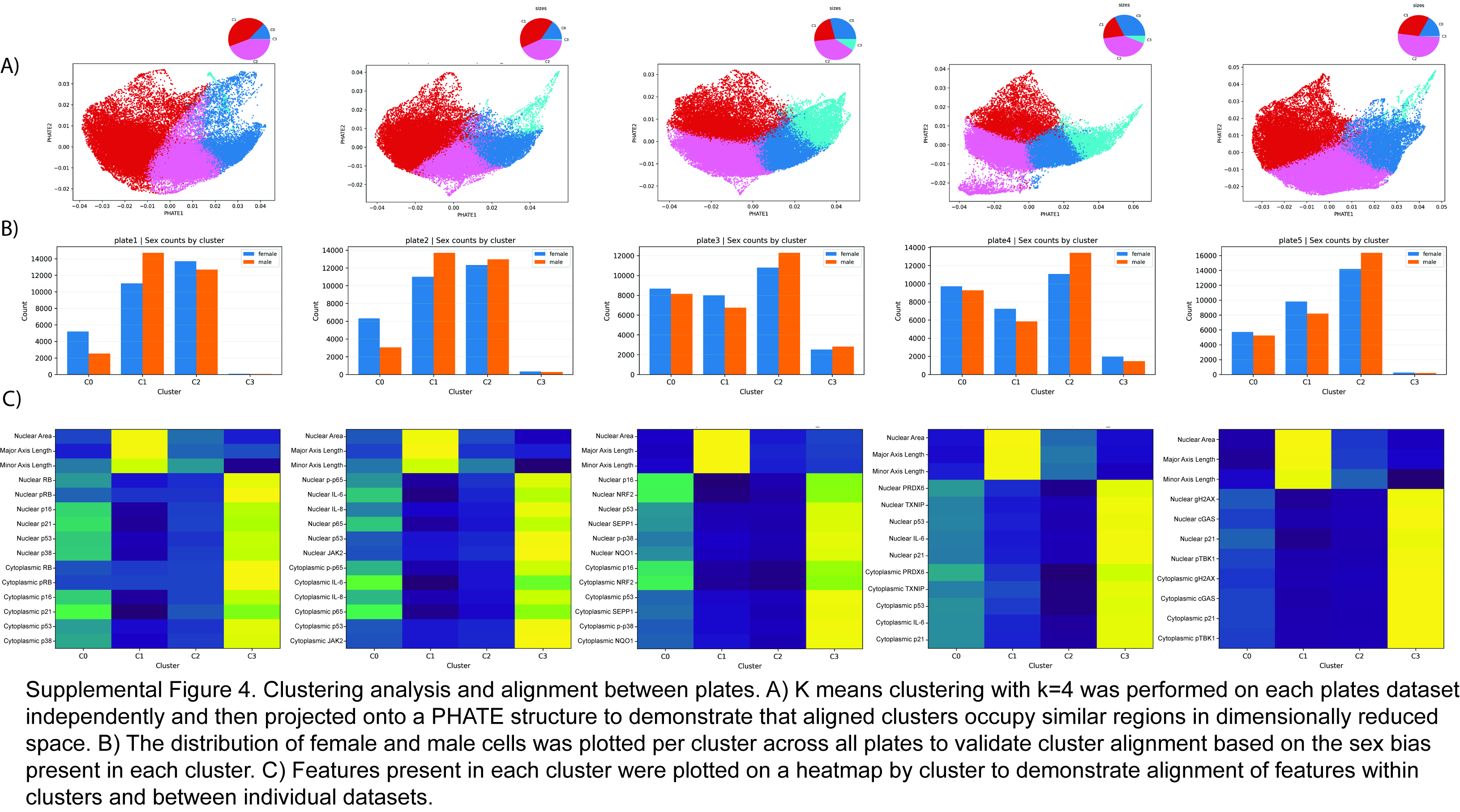

### Supplemental Figure 5

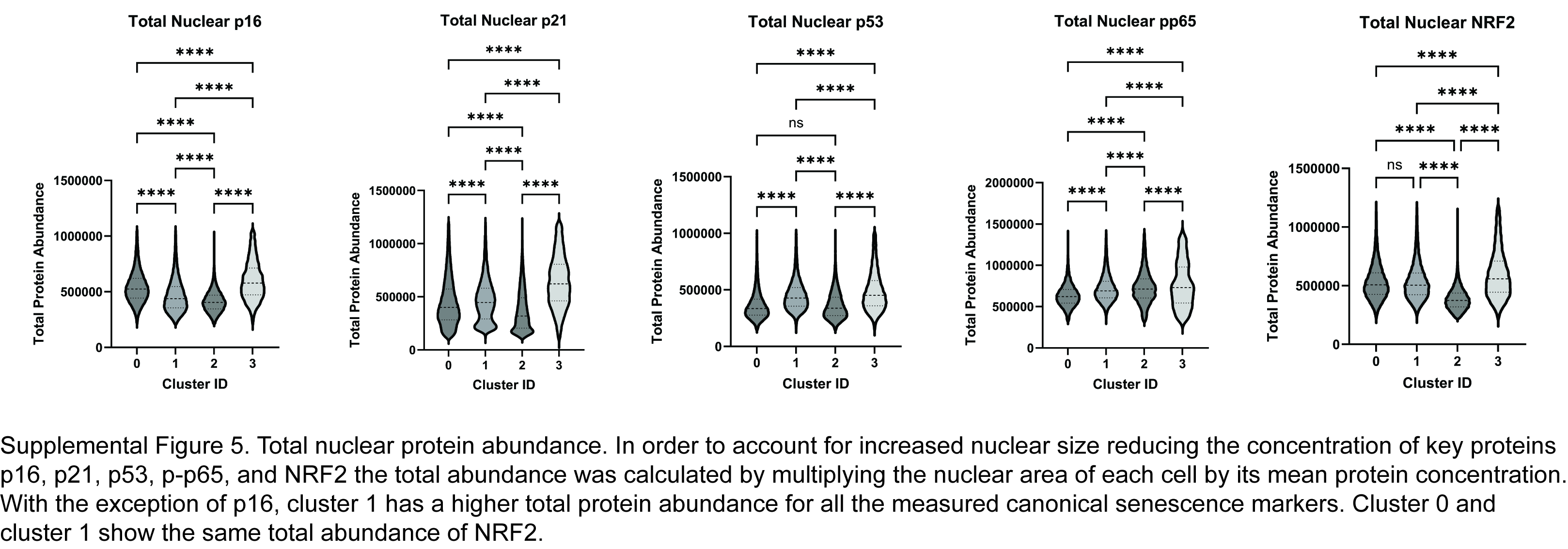

### Supplemental Table 1

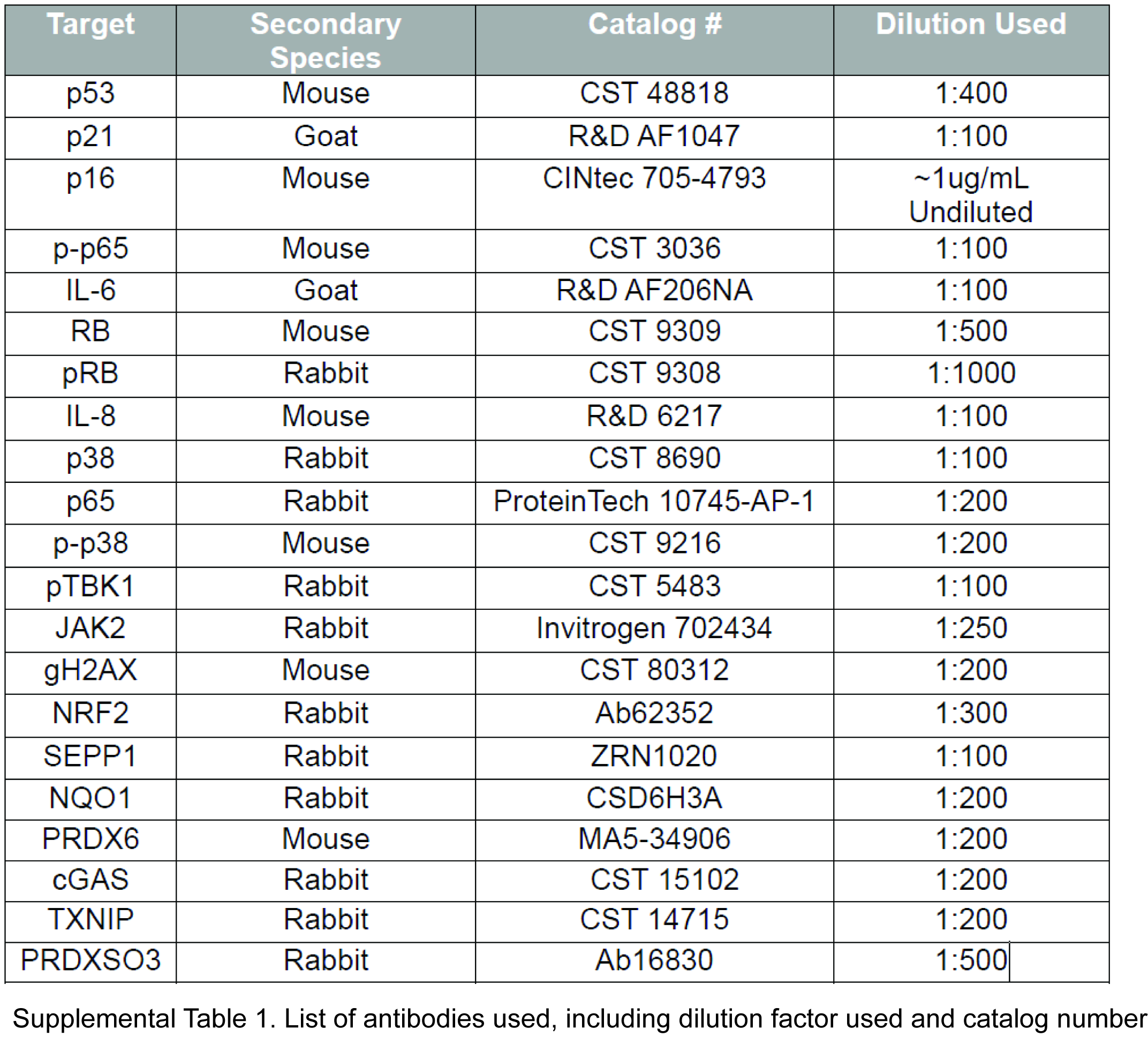

### Supplemental Table 2

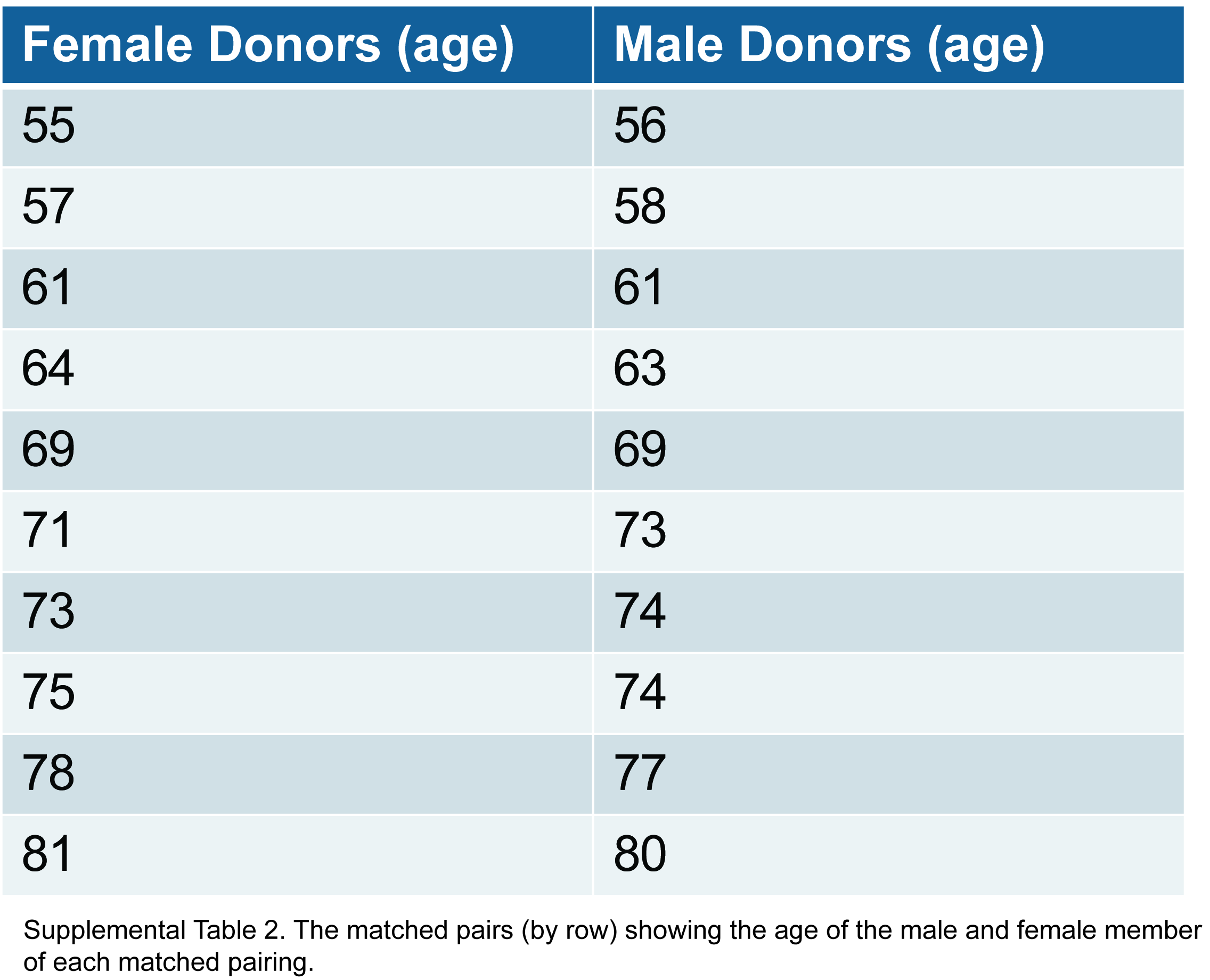

### Supplemental Table 3

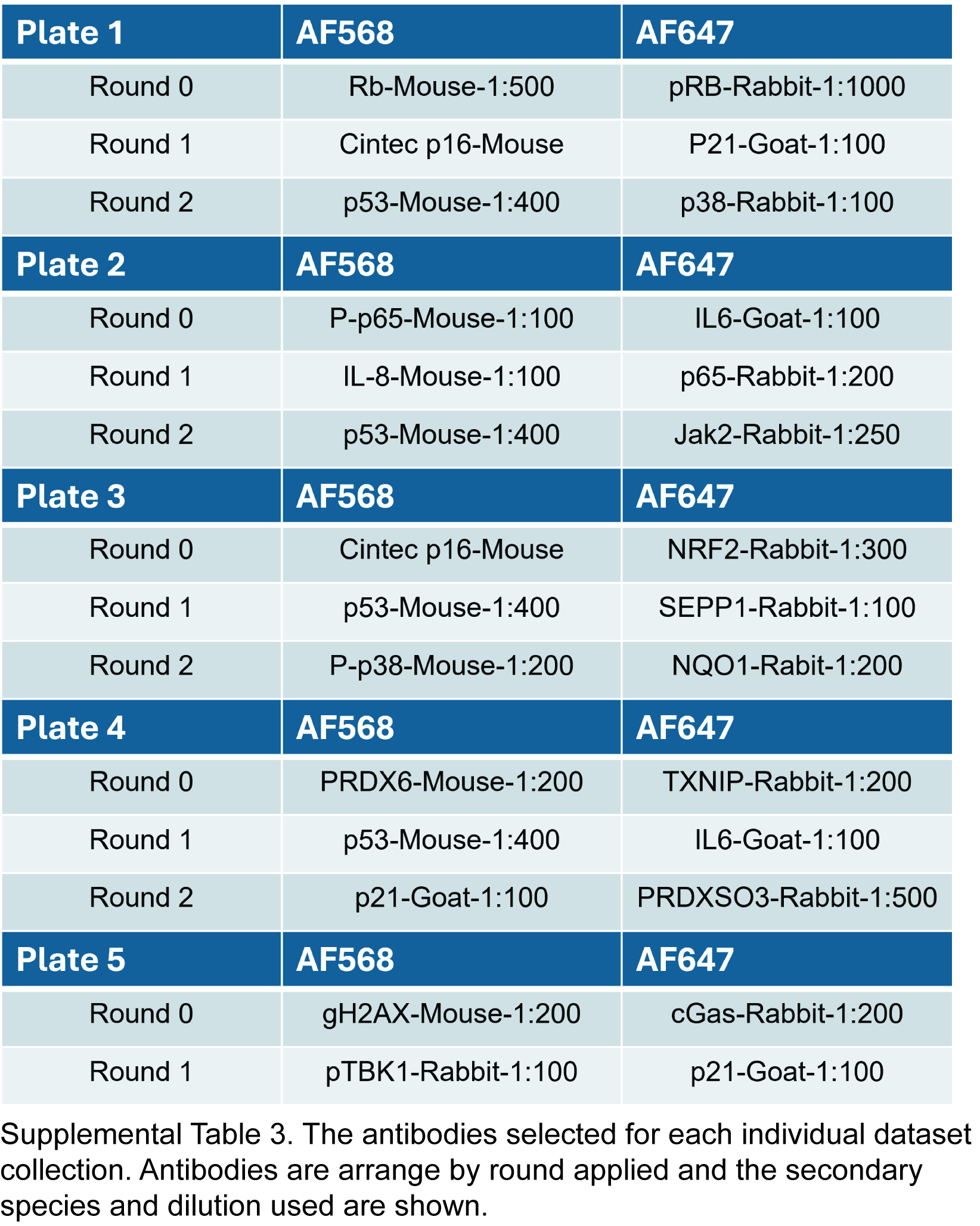
