## Supplemental Table 4 for "Sex differences in senescence burden within human osteoarthritic synovial fibroblasts"

| *Plate* | *Sex* | | *Age* | |
|---|---|---|---|---|
| | Accuracy (RF) | AUC (RF) | R-squared (RF) | RMSE (RF) |
| *Plate1* | 0.85 (0.89) | 0.91 (0.96) | 0.02 (0.24) | 8.10 (7.13) |
| *Plate 2* | 0.74 (0.81) | 0.84 (0.91) | 0.02 (0.22) | 8.08 (7.21) |
| *Plate 3* | 0.68 (0.78) | 0.74 (0.87) | 0.04 (0.23) | 8.01 (7.15) |
| *Plate 4* | 0.77 (0.80) | 0.83 (0.88) | 0.07 (0.19) | 7.89 (7.36) |
| *Plate 5* | 0.80 (0.85) | 0.86 (0.93) | 0.03 (0.30) | 8.06 (6.84) |

Table 4. Model statistics reporting for the two predictive random forest models for sex and age.
